# Characterisation of *Metarhizium majus* (Hypocreales: Clavicipitaceae) isolated from the Western Cape province, South Africa

**DOI:** 10.1101/2020.10.07.329532

**Authors:** Letodi L. Mathulwe, Karin Jacobs, Antoinette P. Malan, Klaus Birkhofer, Matthew F. Addison, Pia Addison

## Abstract

Entomopathogenic fungi (EPF) are important soil-dwelling entomopathogens, which can be used as biocontrol agents against pest insects. During a survey of the orchard soil at an organic farm, the EPF were identified to species level, using both morphological and molecular techniques. The EPF were trapped from soil samples, taken from an apricot orchard, which were baited in the laboratory, using susceptible host insects. The identification of *Metarhizium majus* from South African soil, using both morphological and molecular techniques, is verified. The occurrence of *M. majus* in the South African soil environment had not previously been reported.

## Introduction

Entomopathogenic fungi (EPF), which are cosmopolitan components of the soil microbiota, are commonly isolated from the soil environment, for use as biological control agents to manage a broad range of pest insects [1,2]. The genus *Metarhizium* Sorokin (Ascomycetes, Hypocreales) consists of asexually reproducing EPF species, which are characterised by the production of green conidia on the surfaces of infected insect cadavers, and when they are grown on a growth medium [3]. Species belonging to the genus *Metarhizium* are well-studied entomopathogens, which are widely commercialised. Many products derived from the species are on the market, for use against a wide range of economically-important insect pests [4,5].

Distinguishing between different *Metarhizium* species morphologically is based on their conidial morphology, as using other morphological characteristics is challenging [3]. *Metarhizium* species are mainly identified, and differentiated from each other, using molecular techniques [6]. Two main monophyletic groups fall within the *Metarhizium anisopliae* species complex. The PARB clade consists of *Metarhizium pinghaense* Chen & Guo, *Metarhizium anisopliae sensu stricto*, *Metarhizium robertsii* (Metchnikoff) Sorokin and *Metarhizium brunneum* Petch, while the MGT clade consists of *Metarhizium majus* Johnst., Bisch., Rehner and Humber and *Metarhizium guizhouense* Chen and Guo [3,7]. The MGT species are distinguished from the PARB clade by their relatively large conidia, with *M. majus* having the larger cylindrical conidia relative to *M. guizhouense*, which possess the second largest conidia [7]. *Metarhizium majus* and *M. guizhouense* have been differentiated from each other, based on molecular data, using the translation elongation factor-1 alpha gene (EF-1α) [3,7].

In the current study, additional regarding the morphological and molecular evidence is provided to present the first report on the occurrence of *Metarhizium majus* in South African soil.

## Materials and Methods

### Collection of soil samples and EPF baiting

Soil samples were collected from the orchards surveyed, at a depth of 15 cm, from under the tree canopy on Tierhoek farm (−33°13′19.687″S 19°38′44.281″) (−33,711596; 19,790730), near Robertson in the Western Cape province. After an initial sifting, each soil sample was transferred to 1-L plastic containers, baited with the last-instar larvae of the wax moth *Galleria mellonella* L. (Lepidoptera: Pyralidae) and *Tenebrio molitor* (Coleoptera: Tenebrionidae), kept for 14 days at a room temperature of 25°C [8,9,10]. The soil samples were everted after every 3 days, so as to ensure the penetration of the insect bait into the soil. After every 7 days, the dead insects that showed EPF infection, which was observed in the form of hardening, or overt mycosis of the insect cadaver, were removed from the soil samples. To check the cause of mortality, the dead insects, after first being washed in sterile distilled water, were then dipped in 75% ethanol, followed by them being dipped twice in distilled water. Each dead insect was placed in a Petri dish fitted with moist filter paper. The Petri dishes were then placed in 2-L plastic containers, fitted with paper towels moistened using sterile distilled water, and incubated at room temperature.

Following a further 7 days of incubation, the spores from the surface of the dead insect cuticles were placed on a Sabouraud dextrose agar plate with 1 g of yeast extract (SDAY), supplemented with 200 μl of Penicillin-Streptomycin, so as to prevent bacterial contamination. After the SDAY plates were sealed and incubated at 25°C, they were checked for fungal growth for a period of two weeks. The pathogenicity against insects was verified, using the larvae of the wax moth [11].

### Morphological identification

Temporary slides were prepared by means of trapping spores in a drop of water on a glass slide with a cover slip, secured with glyceel. The size of the conidia was determined, measuring both the length and the width of 30 spores, using a Zeiss Axiolab 5 light microscope, equipped with an Axiocam 208 camera. SEM preparation of spores of different *Metarhizium* species including *M. majus*, *M. robertsii* (GenBank accession number MT378171), *M. pinghaense* (MT895630) and *M. brunneum* (MT380848) were prepared and photographed by the Central Analytical Facility of the Stellenbosch University.

### Molecular identification

For molecular identification, the fungal DNA was extracted from culture plates, using a Zymo research Quick-DNA fungal/bacterial miniprep kit, according to the manufacturer’s protocol. The polymerase chain reaction was conducted, using the KAPA2G ReadyMix PCR kit. Characterisation was based on sequencing of the ITS region and two genes, consisting of the internal transcribed spacer (ITS) (primers ITS1and ITS4), the partial tubulin (BtuB) (primers Bt2a and Bt2b) and the partial EF-1α (primers EF1F and EF2R) [12,13]. The PCR thermocycle conditions accorded with the technique used by [14]. The PCR products were visualised on an agarose gel, using ethidium bromide. The sequences, which were generated by the Central Analytical Facility at Stellenbosch University, were aligned and edited using the CLC main workbench (ver. 8) and blasted on the GenBank database of the National Centre for Biodiversity Information (NCBI) for identification. Fungal cultures were deposited at the Agricultural Research Council (ARC), the Biosystematics Division, Pretoria fungal collection.

### Phylogenetic analyses

Phylogenetic analyses were conducted using the dataset from Rehner and Kepler (2017) [13] and Luz et a. (2019) [12], combining sequences of three loci (ITS, BT, tef1), so as to identify the species concerned. The alignments were done employing ClustalX, using the L-INS-I option. The software package PAUP [15] was used to construct a neighbour-joining phylogenetic tree, using a bootstrap analysis of 1 000 replicates. A Bayesian analysis was run using MrBayes ver. 3.2.6 [16]. The analysis included four parallel runs of 200 000 generations, with a sampling frequency of 200 generations. The posterior probability values were calculated after the initial 25% of the trees were discarded. The fungal isolates used in the current study to construct the phylogenetic trees are listed in Table 1.

**Table 1.**
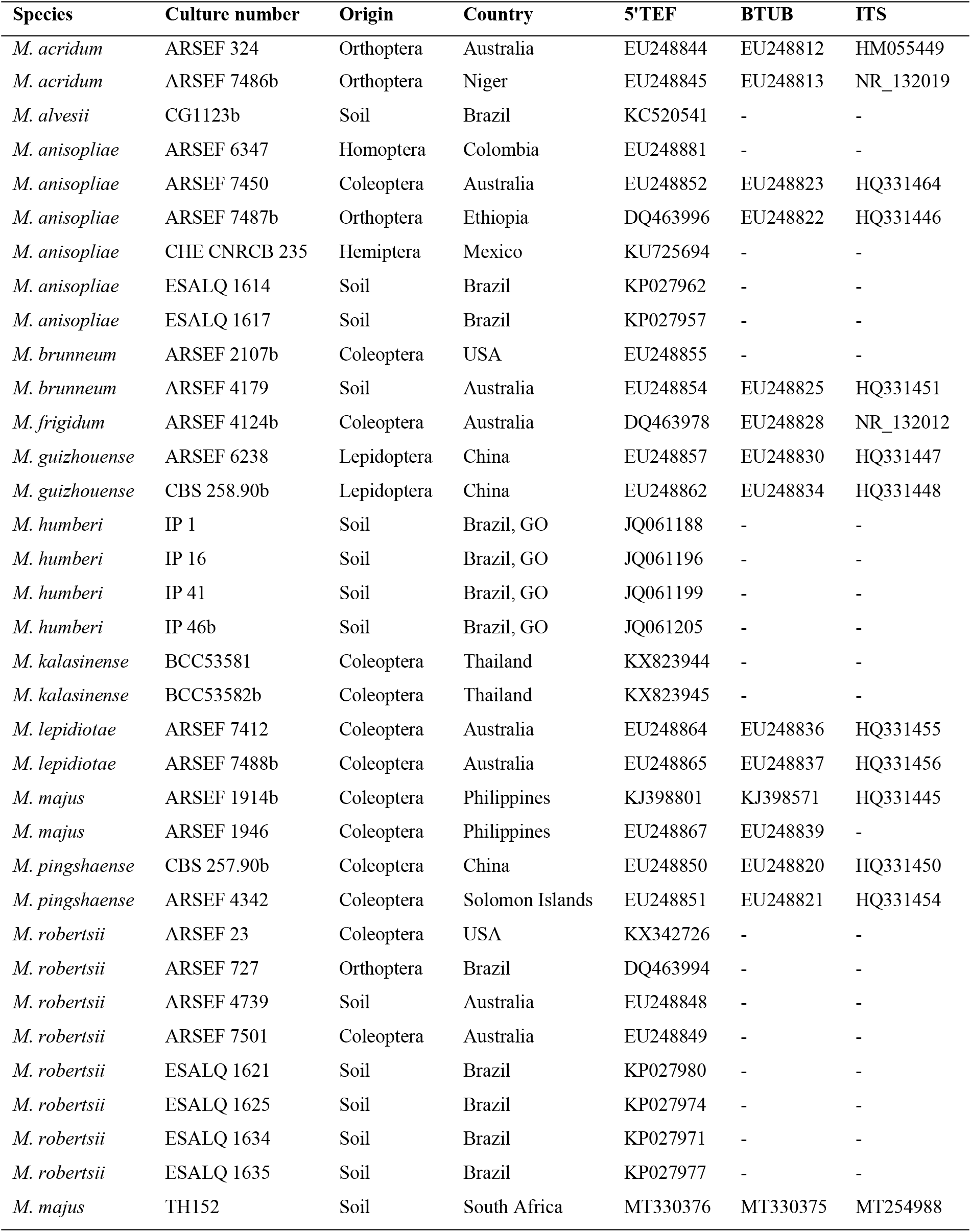
Reference of *Metarhizium* species isolates used in phylogenetic analyses and their culture number, isolation source, country of origin and GenBank accession numbers.

## Results

### Morphological identification

The growth pattern of *M. majus* on SDAY medium was found to typify the genus *Metarhizium* (Fig 1A). The conidia of the mature colonies, which were dark green, formed chains of equal length in clusters (Fig B and C). The conidia, which were oval in shape (n = 30), varied 9.0 (7.5–10.2) μm in length and 4.3 (4.0–4.5) μm in width (Fig 1 D). The conidiophores of *M. majus* were branched, with the apices of the branches bearing from one to several elongated phialides (Fig E and F). SEM pictures of the four different *Metarhizium* species showed no morphological difference in the surface pattern from those of *M. majus* (Fig 1 G and H).

**Fig 1.**
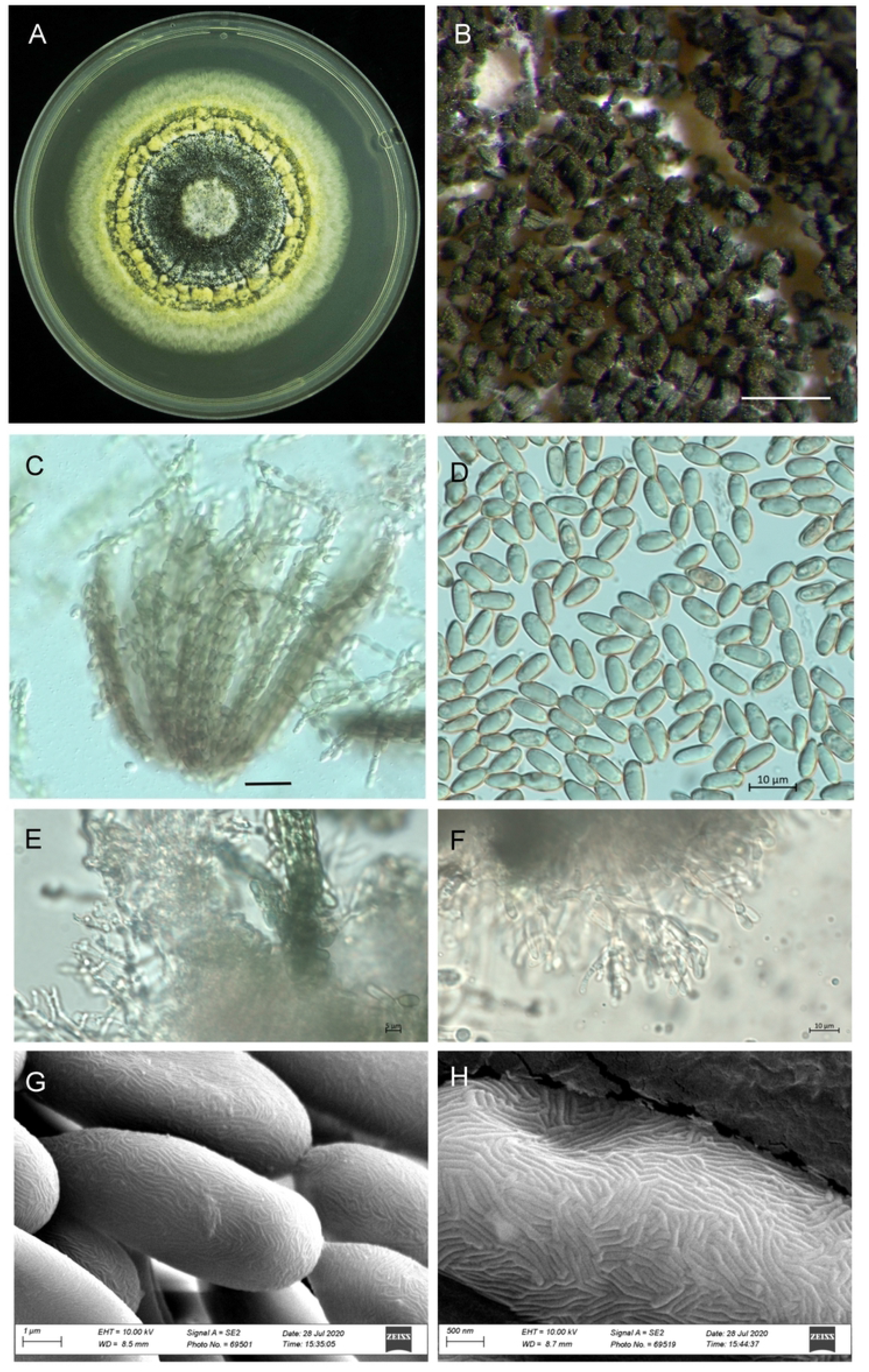
Morphology of *Metarhizium majus* TH152, (A) a three-week-old culture on SDAY medium; (B) spores on older plates; (C) bundles of spore strings of the same length; (D) spore shape and size; (E, F) mature phialides with conidiogenous cells and conidia; (G, H) scanning electron microscope picture showing the surface of the conidia.

### Molecular identification

Sequences generated for the *Metarhizium majus* strain collected from an apricot orchard corresponded to those of the type strains. Using the BLAST function, the ITS region cannot differentiated *M. majus* from *M. anisopliae*, however the elongation factor and the beta-tubulin gene sequences confirm the species to be *M. majus*. Sequences were deposited in the GenBank (ITS: MT254988, EF: MT330376, beta-tubulin: MT330375).

### Phylogenetic analysis

The neighbour-joining phylogeny of the concatenated dataset resulted in a high degree of support for the monophyly of the MGT clades (Fig 2). The *M. guizhouense* was found to form a sister group with *M. majus*, with high percentages of bootstrap support of 87% (Fig 2) and 82% (Fig 3), respectively. The MGT clade forms a sister clade to the PARB clade (Fig 2). The local *M. majus* TH152 isolate, and the two *M. majus* isolates (ARSEF 1914b and ARSEF 1946) collected in the Philippines (Fig 3), grouped in the same clade, with 100% bootstrap confidence.

**Fig 2.**
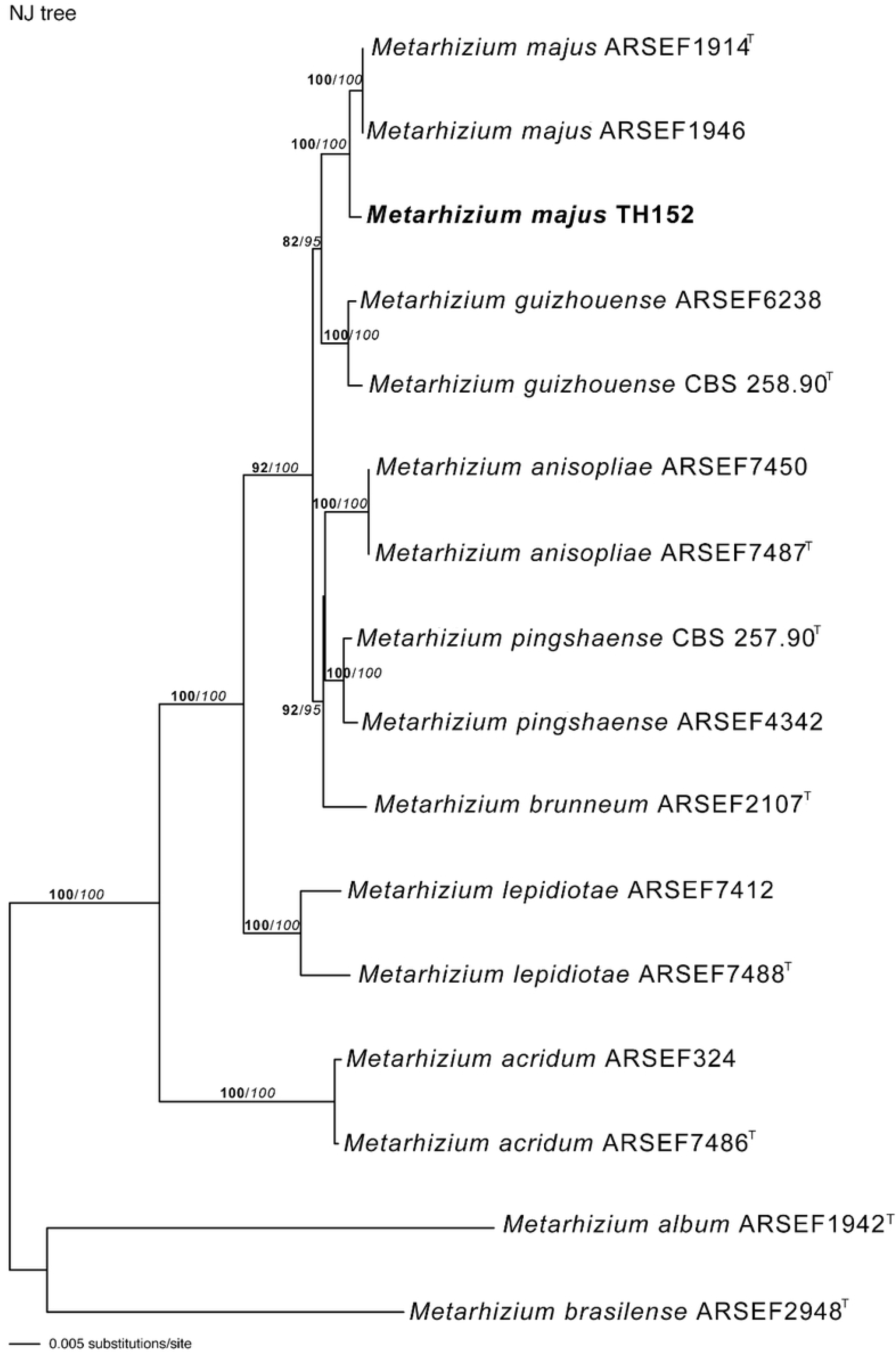
Neighbour joining likelihood phylogenetic tree of *Metarhizium majus* related to the PARB and MGT clades from the analysis of concatenate datasets of 5’intron-rich region of the ITS and the translation elongation factor-1alpha (EF-1α). Bootstrap values are denoted above branches. The tree was rooted, using the sequence from *Metarhizium brasiliense* ARSEF2948b as outgroup and isolates with ^T^ = indicate the type strain.

**Fig 3.**
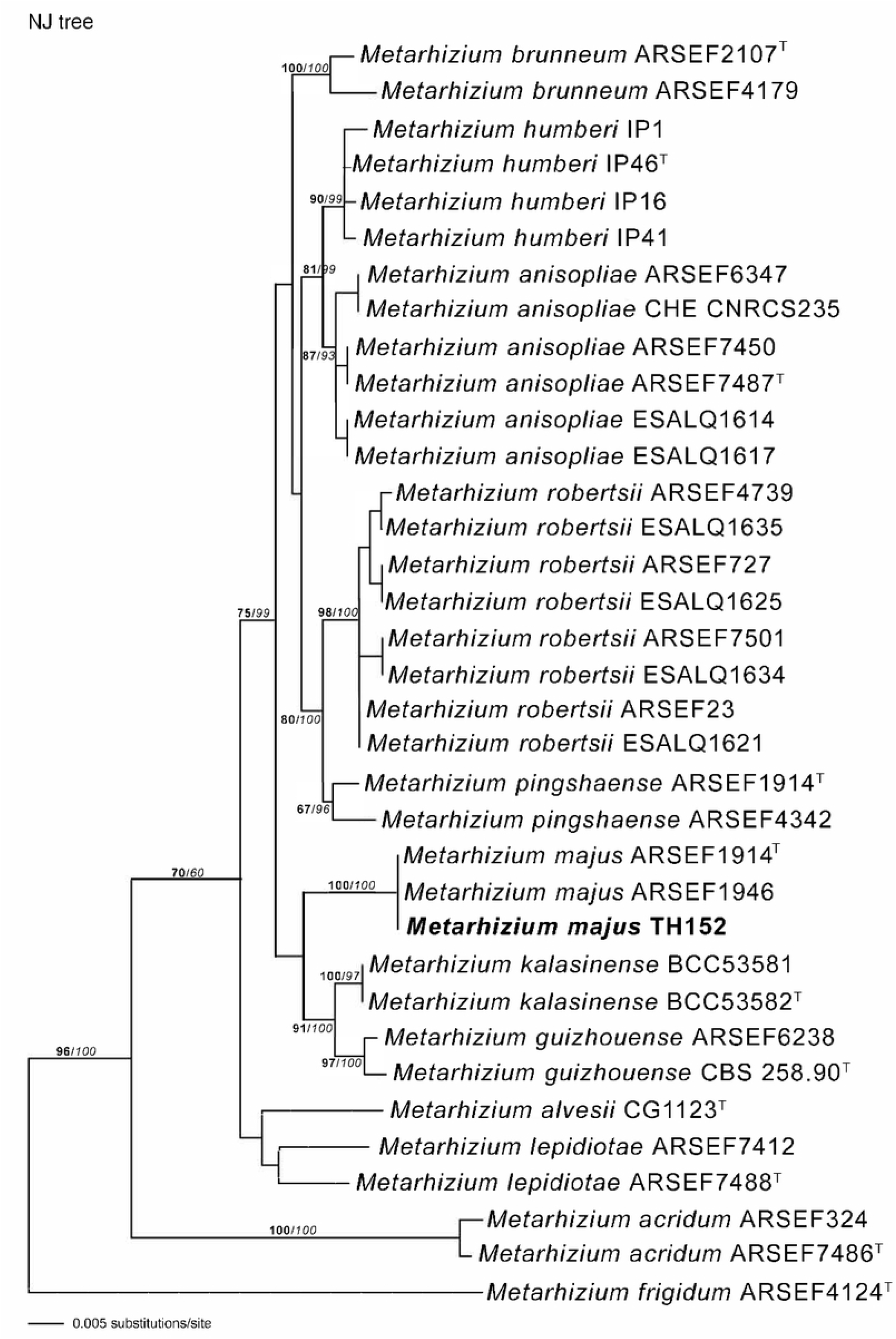
Neighbour-joining phylogenetic tree of *Metarhizium majus* with regards to related species, based on analysis of the 5’intron-rich region of the translation elongation factor 1-alpha (EF-1α) gene sequences. The tree was rooted using the sequence from *Metarhizium frigidum* ARSEF 4124b as the outgroup. Bootstrap values are denoted above branches and isolates with ^T^ = indicate the type strain.

## Discussion

The genus *Metarhizium* consists of a diverse group of entomopathogenic species, having a cosmopolitan distribution and a wide range of insect hosts. *Metarhizium majus* is considered to be an important potential biological control agent for various insect pests. The fungus is deemed to be an effective biological agent, when it is used against *Odoiporus longicollis* Olivier (Coleoptera: Curculionidae), the banana pseudostem weevil, which is a serious pest affecting banana production [18,2]. The EPF is also used to manage *Oryctes rhinoceros* L. (Coleoptera: Scarabaeidae), the coconut rhinoceros beetle, the activities of which result in major crop losses in coconut and palm oil plantations [19,20].

An isolate of *M. majus* was obtained from soil samples, during a survey of EPF in an apricot orchard. The morphological evidence obtained supported the isolate as being *M. majus*, especially in terms of the size of the conidia, which are the largest of all of the *Metarhizium* species. The growth of the hypha is the same for the genus, therefore, being difficult to distinguish from the other species. A previous study indicated that *M. majus* forms one of the species in the group with the largest conidia, ranging from 8.5 to 14.5 μl in length and from 2.5 to 3.0 μl in width, with such a characteristic usually being the only usable morphological difference in the group [3,21]. The surface structure of M. majus was found not to be visually different from four different species of *Metarhizium*.

The presence of *M. majus* in the soil environment has previously been recorded in other countries, like Japan [7] the USA [2], Australia [22] and Denmark [23]. However, it is the first time that the EPF species concerned has been isolated from South African soil, with the current study providing both morphological and molecular evidence of the presence of *M. majus* in such soils. The discovery of the South African *M. majus* isolate adds new information to the body of knowledge regarding soil fungal biodiversity. The presence of *M. majus* in South Africa also increases the number of available local EPF isolates that can be used in agricultural ecosystems for the management of insect pests, as the isolates possess entomopathogenic properties. Its potential as a biocontrol agent, of especially Coleoptera, with be investigated in future studies.

## Acknowledgments

The authors are very grateful to the owner of Tierhoek organic farm. We also wish to thank the all staff and students of the Stellenbosch University participating in the soil sampling.

## Author Contributions

**Conceptualization:** Klaus Birkhofer, Pia Addison, Matthew Addison.

**Formal analysis:** Letodi Mathulwe, Karin Jacobs.

**Funding acquisition:** Pia Addison, Klaus Birkhofer.

**Investigations:** Letodi Mathulwe, Karin Jacobs, Antoinette Malan.

**Project administration:** Pia Addison, Klaus Birkhofer.

**Supervision:** Pia Addison.

**Writing - original draft:** Letodi Mathulwe, Karin Jacobs, Antoinette Malan.

**Writing - review & editing:** Letodi Mathulwe, Karin Jacobs, Antoinette Malan, Klaus Birkhofer, Pia Addison, Matthew Addison

